# Quantitative analysis of inhibitor-induced assembly disruption in human UDP-GlcNAc 2-epimerase using mass photometry

**DOI:** 10.1101/2025.05.10.651030

**Authors:** Nico Boback, Jacob Gorenflos López, Christian P. R. Hackenberger, Santiago Di Lella, Daniel C. Lauster

## Abstract

UDP-GlcNAc 2-epimerase/N-acetylmannosamine kinase (GNE/MNK) is the rate-limiting enzyme in sialic acid biosynthesis and a promising therapeutic target. We applied interferometric scattering microscopy (iSCAM) to investigate GNE oligomerization and its modulation by three small-molecule inhibitors (C5, C13, C15). Substrate binding (UDP-GlcNAc) stabilized tetramer formation by increasing dimer-dimer affinity 120-fold. All inhibitors destabilized tetramers in a concentration-dependent manner, with IC_50_ values in the low micromolar range. Using a modified Cheng-Prusoff equation, IC_50_ values were converted into K_i_ values. Schild analysis was applied to estimate an apparent K_B,app_ value and assess cooperative inhibition effects. Molecular docking confirmed competitive binding for all inhibitors and helped rationalize observed potency trends. While iSCAM has previously been used to study protein assembly, our work demonstrates its applicability for the label-free, quantitative characterization of small-molecule inhibitors affecting protein oligomerization. These findings provide a foundation for further mechanistic studies and underscore the potential of iSCAM in drug-target interaction profiling.

## INTRODUCTION

The *de novo* synthesis of sialic acid (SA), a key sugar involved in cellular processes such as signaling, adhesion, and immunity,^1-3^ is governed by the bifunctional enzyme UDP-*N*-acetylglucosamine 2-epimerase/*N*-acetylmannosamine kinase (GNE/MNK). As the rate-limiting enzyme in SA biosynthesis, GNE/MNK converts uridine diphosphate *N*-acetylglucosamine (UDP-GlcNAc), a product of the hexosamine biosynthetic pathway, into *N*-acetylmannosamine (ManNAc), which is subsequently phosphorylated under ATP consumption to form ManNAc-6-phosphate (ManNAc-P_6_). Two additional conversion steps, catalyzed by sialic acid synthase (SAS) and sialic acid phosphatase (SAP), yield the final product, Neu5Ac, which can be further converted by CMP-Neu5Ac synthetase into the feedback inhibitor CMP-Neu5Ac.^4-6^ Individual GNE subunits primarily form enzymatically inactive homodimers, but the presence of UDP-GlcNAc promotes formation of an active homotetrameric state.^4,5,7^ However, even in its active form, GNE exhibits low catalytic efficiency (k_cat_ = 11.8 ± 2.0 s^-1^, and K_M_ = 33.1 ± 4.2 µM). In contrast to the substrate, the feedback inhibitor CMP-Neu5Ac stabilizes an inactive tetrameric conformation by binding at the dimer-dimer interface, thereby downregulating enzyme activity.^4^ This assembly-state-dependent regulation of GNE is a central focus of the present study.

The crystal structure of GNE, derived from a complex with UDP (substrate component) and CMP-Neu5Ac (feedback inhibitor), reveals that each GNE monomer (∼43 kDa) contains a substrate-binding site within its core. UDP interacts at this site through multiple salt bridges and hydrogen bonds (**Figure 1A**). GNE forms dimers through non-covalent interactions involving hydrophobic side chains from helices α3, α4 and α5 at the *N*-terminal region. These dimers further associate into tetramers *via* a 2+2 binding configuration (**Figure 1B**), stabilized by a hydrophobic core at helix α3 near the tetramer interface and multiple salt bridges.^4^ The MNK subunit connects to the *C*-terminus of the GNE subunit (**Figure 1A**, yellow), establishing the full-length protein “Master Regulator”, forming homodimers in absence or homotetramers in presence of substrate or feedback inhibitor.^4,5,8^ The GNE subunits *C*-terminal connecting region exhibits high flexibility, facilitated by a hinge region involving Gly182 and Asp187.^8^ A full-length structural model suggests that the active sites of GNE and MNK are positioned in close proximity, potentially forming an internal epimerase-kinase channel to enable efficient and protected transfer of intermediates.^4,5^

**Figure 1.**
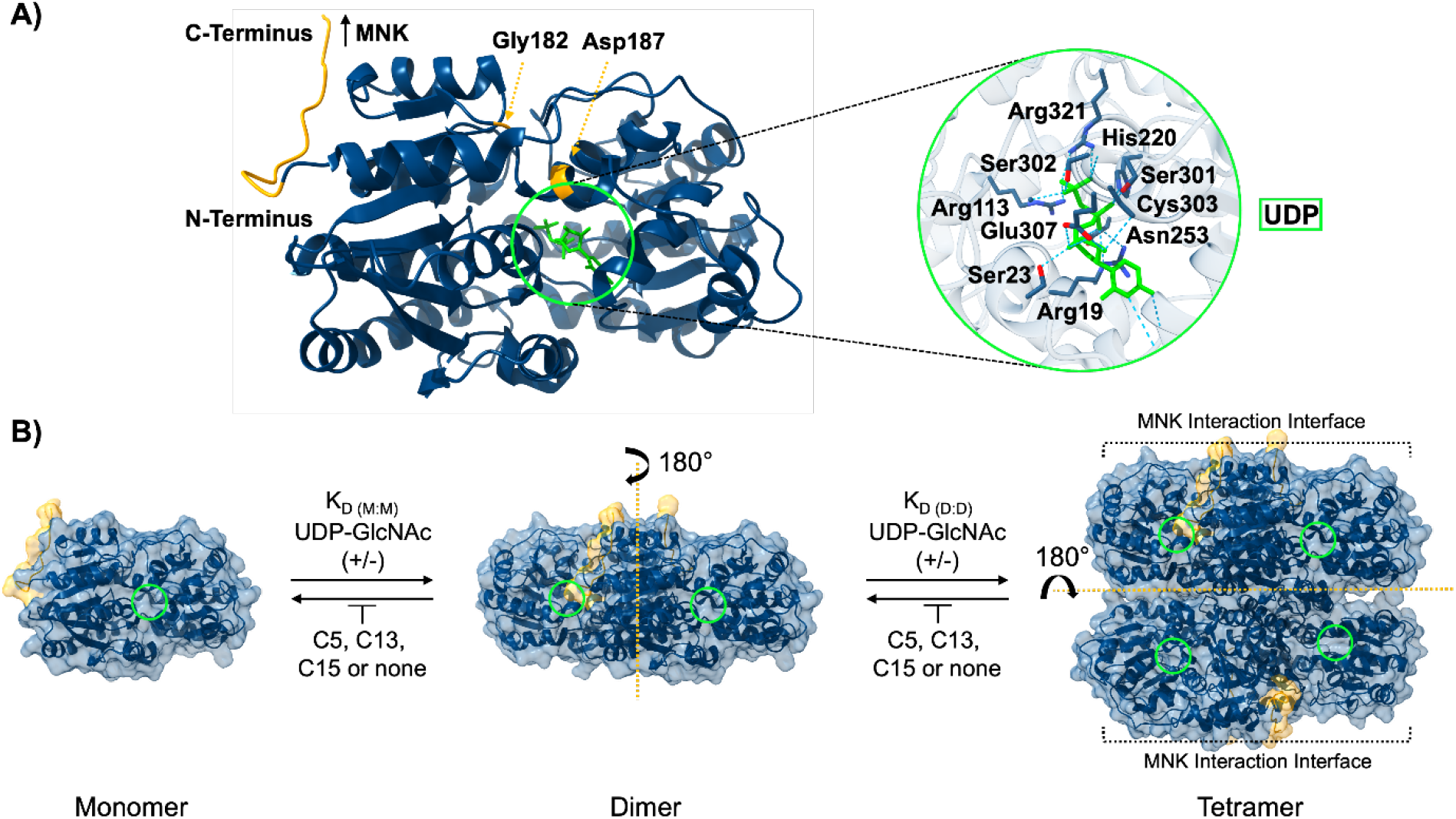
Structural insights into GNE-substrate binding and tetramer assembly dynamics in the presence of substrate or inhibitors. A) Crystal structure of monomeric GNE bound to UDP (green), highlighting key protein residues involved in substrate interaction. B) GNE assembly equilibria between monomer-dimer (K_D(M:M)_), and dimer-tetramer (K_D(D:D)_) states, characterized by distinct dissociation constants in the presence or absence of UDP-GlcNAc and inhibitors (C5, C13, C15). The substrate binding site is indicated by a green circle. The *C*-terminal region connecting GNE and MNK subunits in the full-length protein and key residues increasing linker flexibility are highlighted in yellow. The monomeric structure and protein assemblies of GNE are derived from the protein data bank (PDB, ID: 4zht).^4^ Structural visualizations were generated using ChimeraX-1.9.

Targeting the GNE/MNK enzyme – either as a whole or through its individual protein subunits – offers a potential strategy for therapeutic intervention. This approach has been demonstrated in the modulation of SA densities on cell surfaces,^9,10^ impacting cellular signaling,^1,2^ and host-cell susceptibility to pathogens, including influenza A virus.^1,3^ In the context of influenza A virus inhibition, reducing sialic acid densities on the respiratory epithelium has proven effective. Strategies such as CRISPR-based GNE knockouts^11^ and sialidase fusion proteins^12^ have successfully inhibited viral infection. However, unlike permanent gene knockouts, small-molecule inhibitors of GNE could provide a reversible and controlled alternative for modulating SA biosynthesis, offering greater therapeutic flexibility.

Previous enzyme activity assays from our group established that GNE’s quaternary structure directly influences its enzymatic activity. UDP-GlcNAc binding promotes tetramer formation, while small-molecule inhibitors disrupt oligomerization by interfering with substrate binding.^7^ Additionally, the feedback inhibitor CMP-Neu5Ac was shown to stabilize the tetrameric, inactive state *via* allosteric modulation, altering UDP-GlcNAc affinity at the substrate binding pocket.^4^

A high-throughput screening of 68,640 small-molecule compounds identified three potent, non-carbohydrate-based UDP-GlcNAc 2-epimerase inhibitors: C5, C13 and C15. These inhibitors exhibited high potency and cell culture medium stability, with C5 derived from the purine base xanthine scaffold, while C13 and C15 share a pyrimidinone core, structurally similar to UDP-GlcNAc, which is linked *via* a vinyl to an aromatic system. Further characterization of these inhibitors revealed distinct destabilizing effects on GNE assembly. Hydrogen-deuterium exchange mass spectrometry (HDX-MS) results suggested that all three inhibitors compete with UDP-GlcNAc for the substrate binding site, leading to protein disassembly. However, C13 and C15 also induced unique conformational changes, distinct from those caused by C5 or the substrate, enhancing negative cooperativity in protein assembly and thus inhibitor potency. Qualitative analysis confirmed a progressive increase in inhibitory potency from C5 to C13 to C15. Despite these insights, the precise molecular mechanisms underlying GNE assembly inhibition remain only partially understood, as different inhibitors may disrupt oligomerization through diverse pathways.^7^

To further explore the effects of these inhibitors, we previously employed interferometric scattering microscopy (iSCAM), also known as mass photometry (MP), to study the stability of GNE tetramers and dimers at a single inhibitor concentration (10 µM) in the presence or absence of UDP-GlcNAc (10 µM).^7^ This technique detects mass-dependent signals from individual proteins or protein assemblies by measuring interferometric light scattering at a glass-solution interface.^13,14^ Our findings demonstrated that C5, C13 and C15 all disrupted tetrameric GNE, while C13 and C15 also destabilized dimeric assemblies, a property not observed for C5. Consistent with enzyme activity assays, C13 and C15 exhibited higher potency than C5, suggesting a stronger influence on GNE disassembly.^7^ However, the underlying differences in potency and inhibition mechanisms remain incompletely understood.

Building upon our initial findings, the present study was designed to systematically characterize the inhibition mechanisms of C5, C13 and C15 using iSCAM. Although this technique has been applied to various biological systems, including bovine serum albumin (BSA),^14^ plant-derived 2-cysteine peroxiredoxins,^15^ transmembrane transport channels (KcsA, OmpF), membrane oxidoreductases,^16^ antibodies (IgG),^17,18^ SARS-CoV-2 spike proteins,^19^ and surfactant proteins A and D,^20^ studies examining the inhibition of protein assembly using iSCAM remain limited.^7,21^ Compared to conventional methods such as gel electrophoresis,^6^ analytic size exclusion chromatography (aSEC), and hydrogen-deuterium exchange mass spectrometry (HDX-MS),^7^ iSCAM offers key advantages, including low sample consumption, label free single-molecule detection, and a broad molecular mass range (40 kDa to ∼10 MDa) with high precision (1 kDa).^14,15,22^

This study serves a dual purpose. First, it provides a quantitative characterization of GNE assembly and inhibition. To this end, we systematically investigated GNE assembly equilibria in the presence or absence of UDP-GlcNAc, determining monomer-monomer (M:M) and dimer-dimer (D:D) binding affinities (K_D_ values) as indicators of GNE complex stability. The effects of the inhibitors, C5, C13, and C15, on these equilibria were quantitatively assessed using iSCAM, yielding IC_50_ values. These values were then converted to K_i_ values under the assumption of mixed competitive and allosteric inhibition – suggested by experimental results – using a modified Cheng-Prusoff equation (**Equation S4**) that accounts for allosteric effects induced by orthosteric inhibitors.

To demonstrate the applicability of our approach, we conducted a representative Schild analysis for inhibitor C15, applying modified Schild equations (**Equation S5, S6**) that incorporate both allosteric and competitive interactions observed in the inhibited GNE system. The resulting Schild plot and function (**Figure 4, S4)** allowed visualization of inhibition dynamics and quantification of the allosteric effect induced by the antagonist (inhibitor). While Schild analysis is widely used to evaluate antagonist potency and binding mechanisms, especially in competitive systems, its assumptions may not fully capture complex, cooperative behaviors.^23^ To address this, we additionally employed the Operational Model of Allosterically Modulated Agonism (OMAM) to fit substrate response data in the presence of C15.^24^ This model enabled us to extract mechanistic parameters such as efficacy (**τ**), the substrate (K_S_) and inhibitor (K_B_) affinity to GNE, and cooperativity factors (α, β), thereby offering a more detailed and quantitative description of the allosteric inhibition observed. Finally, computational docking analyses provided atomistic insights into inhibitor binding, further supporting the mechanistic conclusions derived from both Schild and OMAM analyses.

Second, this study establishes iSCAM as a powerful method for investigating protein assembly inhibition. Its single-molecule, label-free format enabled us to quantify how small molecules modulate GNE oligomerization. By analyzing monomer-monomer and dimer-dimer binding affinities (K_D_ values) and Hill coefficients, we characterized both binding dynamics and allosteric effects. IC_50_ values from concentration-dependent inhibition curves were translated into K_i_ values, and a modified Schild and OMAM analysis for inhibitor C15 illustrated the applicability of the approach to assembly-based inhibition mechanisms.

## RESULTS

We systematically investigated GNE assembly dynamics under controlled conditions using recombinant His_6_-tagged GNE, which exhibited an observed molecular mass of 46.4 kDa. The protein was prepared following previously published protocols by Gorenflos López and colleagues.^6,7^ All experiments were conducted at 20 °C in DPBS (pH 7.4), with samples maintained in solution. To ensure thermodynamic equilibrium, each sample was incubated for 30 minutes before measurements.

### GNE assembly equilibrium in the absence of UDP-GlcNAc

To quantify GNE assembly equilibria in the absence of substrate, we performed mass photometry experiments on a serial dilution of GNE in DPBS, with protein concentrations ranging from 50 nM to 50 µM (**Figure 2A**). Throughout the experiment, we identified three distinct molecular assemblies at 50 kDa, 90 kDa and 180 kDa, corresponding to monomers, dimers, and tetramers, respectively (**Figure 2B**).

**Figure 2.**
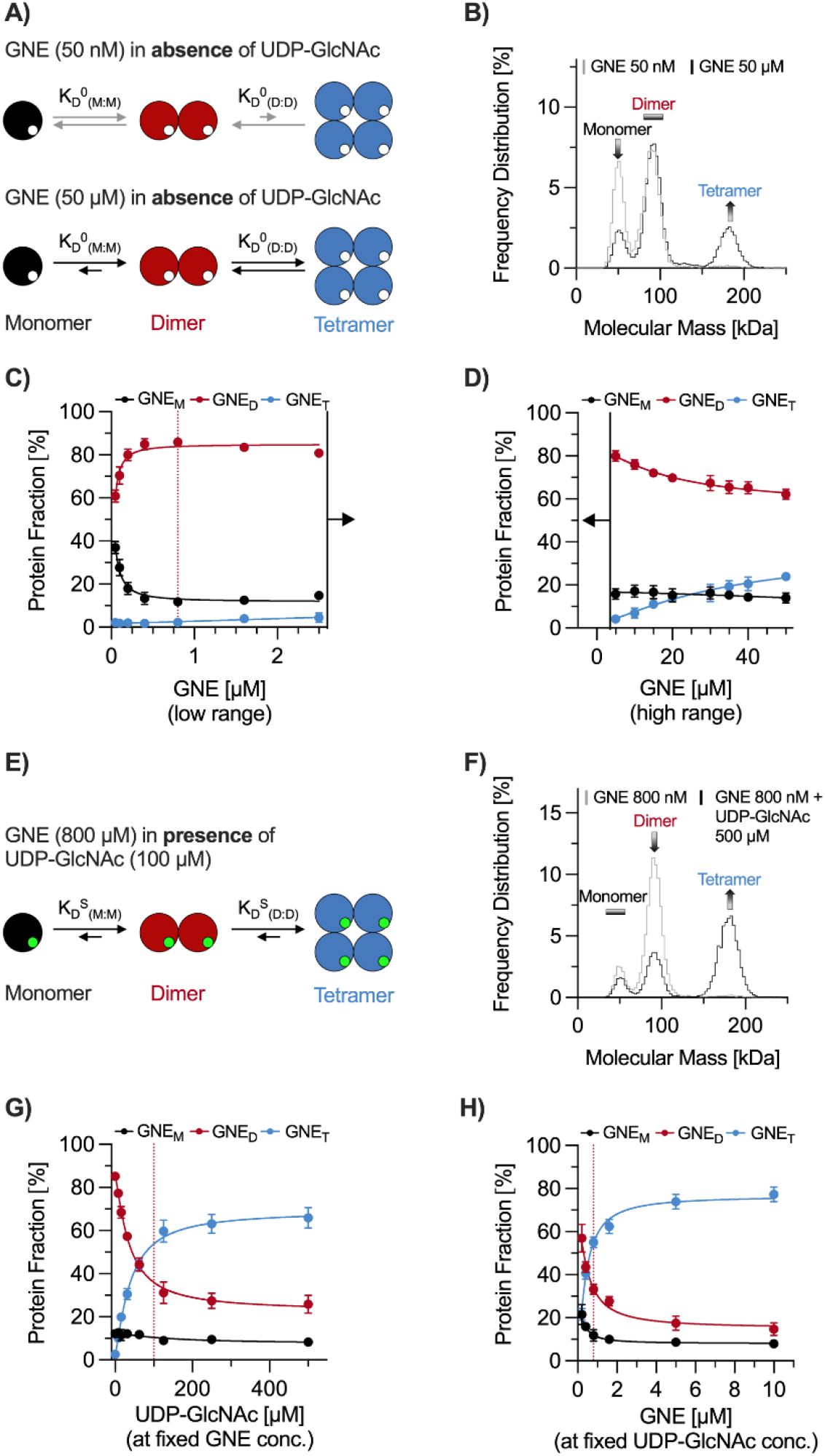
Assessment of GNE assembly dynamics using mass photometry. (A) Schematic representation of monomer-dimer (M:D) and dimer-tetramer (D:T) equilibria in the absence of UDP-GlcNAc (0 indicates the absence of substrate), leading to affinity constants (K_D_^0^). Large circles represent GNE, while small empty circles denote unoccupied substrate-binding sites. At a low GNE concentration (50 nM) the assembly equilibrium is shifted to monomers (grey arrows), while at a high GNE concentration (50 µM) the assembly equilibrium is shifted to tetramers (black arrows). B) iSCAM histograms of GNE at 50 µM (black) and 50 nM (grey) corresponding to the scheme in A). C and D) Effect of GNE concentration on its assembly distribution, expressed as protein fraction of each subunit (M = monomer, D = dimer, T = tetramer) relative to the total protein concentration, shown for low (C) and high (D) protein concentration regimes. The dotted red line marks the minimal GNE concentration (800 nM) required to reach a plateau in protein assembly, which was fixed for subsequent experiments. E) Schematic representation of UDP-GlcNAc-dependent (S indicates the presence of substrate) GNE M:D and D:T equilibria (indicated by arrows), leading to affinity values (K_D_^S^). Green-filled circles indicate UDP-GlcNAc occupying the substrate-binding pocket. F) iSCAM histograms of GNE (800 nM) in the absence (grey) and presence (black) of UDP-GlcNAc (100 µM), corresponding to the scheme in E). G) and H) Effect of UDP-GlcNAc concentration on GNE assembly distribution. G) Substrate titration against a fixed GNE (800 nM) and H) GNE titration against a fixed substrate concentration (100 µM). In G), the dotted red line indicates the minimal UDP-GlcNAc concentration required to reach maximum tetramer formation. In H), it marks the fixed GNE concentration (800 nM) used in subsequent experiments. All histograms include at least 8000 counts. The “Frequency Distribution” represents the percentage of detected counts corresponding to a specific mass relative to the total number of detected counts. Error bars indicate the standard deviation from three independent experiments, each performed in technical triplicate. Graphs are color-coded to indicate GNE monomers (M, black), dimers (D, red), and tetramers (T, blue). Data fitting was performed using a Hill-based logistic fit (**Equation S1**) adapted from the Kukura lab,^13^ with a Hill-coefficient n = 1.3 determined in (G).

Comparing these measured masses to the apparent monomer mass of 46.4 kDa, previously reported in our studies,^6,7^ the iSCAM results demonstrate 93% mass accuracy for the monomer and 97% for the dimer and tetramer using our experimental setup and mass standards. Between measurements, mass deviations remained within 1 - 3 kDa, indicating high reproducibility. As GNE concentration increased, we observed a shift in the protein assembly ratio towards higher molecular mass populations from 17:28:1 (M:D:T) at 50 nM to 4:18:7 at 50 µM (**Figure 2A, B**). Up to a GNE concentration of 800 nM, the dimer fraction increased, while monomer fractions decreased, reaching a stable multimer distribution over an extended concentration range. However, at protein concentrations above 5 µM, we observed a concomitant rise in tetramer formation and a decrease in dimer levels (**Figures 2C, D**). To determine protein affinities (K_D_) between monomers and dimers, protein fraction data ([GNE_M/D/T_]/[GNE_total_]) was plotted against the concentration of free monomer (GNE_M,free_) or free dimer (GNE_D,free_) (**Figure S1A, B**). The data were fitted using a Hill-based logistic model (**Equation S1**),^13^ derived from the law of mass action, with an Hill-coefficient of n = 1.3, determined in the following substrate titration experiments. This analysis enabled us to determine the binding affinity constants (K_D_), which represent the half-maximal conversion of monomer into dimer (K_D_^0^_(M:M)_) or dimer into tetramer (K_D_^0^_(M:M)_). We determined a monomer-monomer (M:M) interaction K_D_^0^_(M:M)_ of 10.9 ± 1.0 nM, while the dimer-dimer (D:D) interaction exhibited a K_D_^0^_(D:D)_ of 23.5 ± 4.8 µM. These values indicate that monomers associate more strongly to form dimers compared to dimers assembling into tetramers, highlighting a greater stability of the dimeric state in the absence of substate.

### Substrate effect on dimer and tetramer stability of GNE

To examine the impact of UDP-GlcNAc (substrate) on GNE subunit affinities, we conducted a series of titration experiments (**Figure 2E**). In the first part of the experiment, UDP-GlcNAc was titrated against a fixed GNE concentration (800 nM), which represents the minimum concentration required to establish a stable assembly equilibrium based on our previous dilution studies. As the UDP-GlcNAc concentration increased, we observed a progressive shift toward tetramer formation, which is clearly reflected in the mass photometry spectra (**Figure 2F**). A plateau in tetramer formation was reached at 100 µM UDP-GlcNAc, which was subsequently used as the standard concentration for all following experiments (**Figure 2G**). From the slope of the fitted curves (**Equation S1**),^13^ we determined a Hill coefficient of n = 1.3, indicating a strong positive cooperativity. This value was fixed for fitting the data across both, the previous dilution and subsequent inhibition experiments. We further determined, from the tetramer curve, a moderate substrate affinity (K_S_) of 34.6 ± 3.0 µM using **Equation S2**, which is close to the K_M_ (33.1 ± 4.2 µM) of GNE.^4^

In the second part of the experiment, we performed the reverse titration, in which GNE was titrated against a fixed concentration of UDP-GlcNAc (100 µM) (**Figure 2H**). Similar to the first titration, an increasing GNE concentration led to a continuous shift towards tetramer assembly.

To quantify the intermolecular affinities (K_D_) between monomers and dimers, we plotted the protein fraction data against the concentration of free monomer (GNE_M,free_) or free dimer (GNE_D,free_) (**Figure S1C, D**) and fitted the data using a Hill-based logistic model (**Equation S1**).^13^ The fitted curves yielded K_D_ values of 55.8 ± 5.2 nM for M:M interactions and 196.2 ± 18.2 nM for D:D interactions, indicating that UDP-GlcNAc promotes GNE oligomerization, by significantly enhancing the stability of tetramers.

### GNE Assembly Inhibition Studies

Building on the stoichiometric distribution of GNE assemblies, as defined by the K_D_ values for dimers and tetramers under specific conditions, we further investigated the potency of three non-carbohydrate-based inhibitors (C5, C13, and C15) in destabilizing GNE dimers and tetramers, determining their respective IC_50_ values. To assess inhibition, we performed titration experiments, in which varying concentrations of inhibitors were tested against a constant GNE concentration (800 nM), both in the presence or absence of UDP-GlcNAc (100 µM).

In the presence of UDP-GlcNAc, all inhibitors effectively destabilized tetramers (**Figure 3A-C**), as further illustrated schematically in **Figure 3D**. However, C13 and C15 also exhibited an additional destabilizing effect on dimers (**Figure 3E, F**). Notably, inhibition by C13 and C15 followed a sequential destabilization pattern, initially targeting tetramers, followed by dimers, allowing for the differentiation between low (**Figure 3B, C**) and high (**Figure 3E, F**) inhibitor concentration regimes.

**Figure 3.**
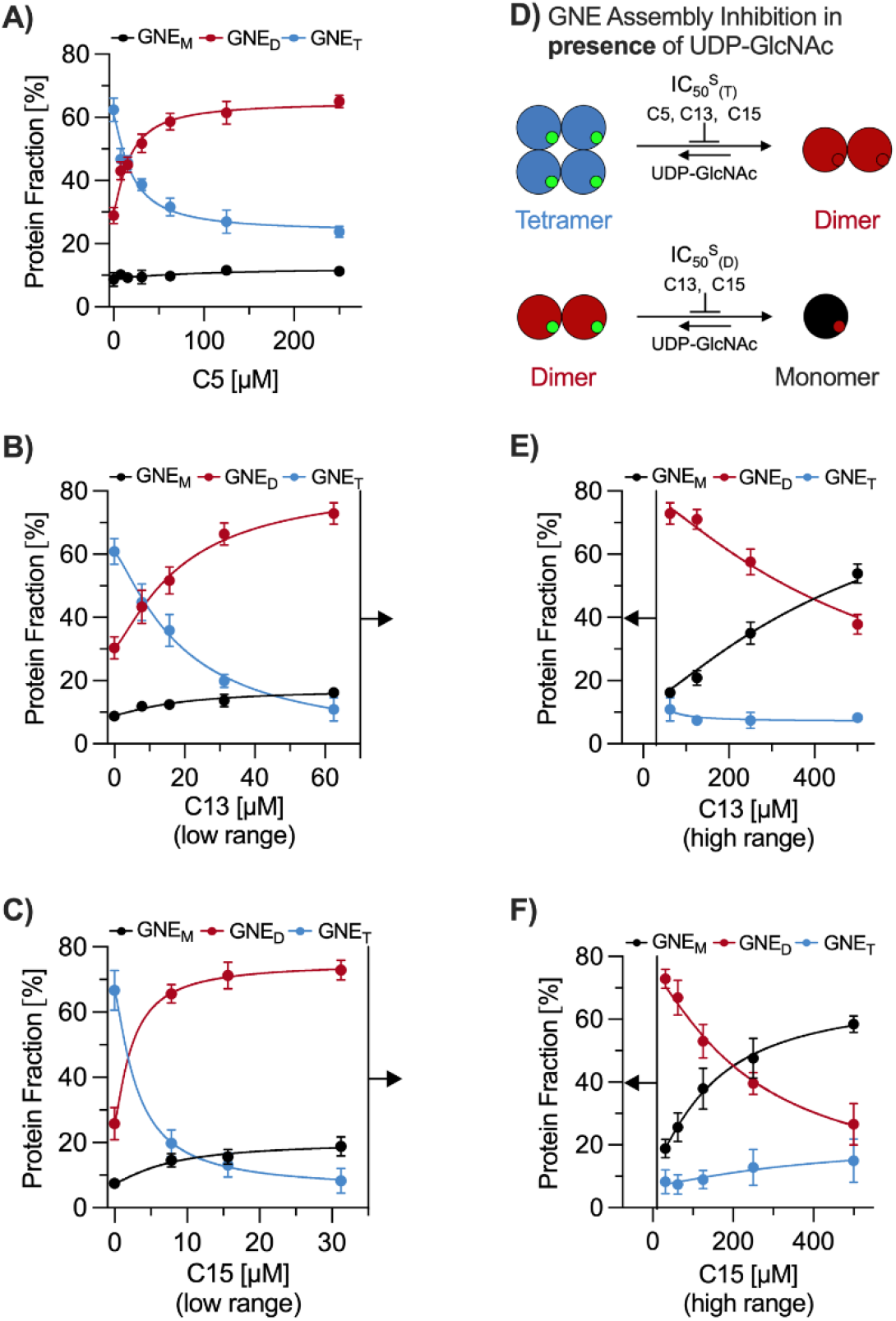
Inhibition of GNE assembly by C5, C13, and C15 using mass photometry. The concentration-dependent effects of inhibitors on the GNE (800 nM) tetramer and dimer stability were assessed using mass photometry. The monomer (M), dimer (D) or tetramer (T) fraction, expressed as a percentage of the total protein concentration, is shown for A) C5, B) C13 and C) C15 at low inhibitor concentrations. D) Schematic representation of the inhibition model, illustrating the equilibrium shift from tetramer (T, blue) to dimer (D, red) and dimer to monomer (M, black) upon inhibitor binding. This equilibrium was analyzed in the presence of 100 µM UDP-GlcNAc, denoted by superscript S, to determine inhibitor-specific IC_50_ values for tetramer (IC_50_^S^_(T)_) and dimer (IC_50_^S^_(D)_) destabilization. The large colored circles represent GNE protein assemblies, while small green and red filled circles indicate the substrate binding pocket occupied by UDP-GlcNAc (green) or inhibitor (red). The inhibitor concentration-dependent effects on GNE assembly distribution were also assessed at higher concentrations for E) C13 and F) C15 extending the lower concentration range shown in B) and C). Error bars represent the standard deviation from three independent experiments, each performed in technical triplicate. Data were fitted using a Hill-based logistic function (**Equation S3**), with a Hill-coefficient n = 1.3, determined in **Figure 2G**.

In the absence of UDP-GlcNAc, C13 and C15 significantly disrupted the dimeric assembly, whereas C5 had a minimal effect (**Figure S3**). By plotting protein fraction data against the inhibitor concentration, we determined IC_50_ values (**Table 1, S1**) using a Hill-based logistic model (**Equation S3**). We further translated the IC_50_ values into assay-independent K_i_ values, under the assumption of a mixed competitive and allosteric inhibition, using a Cheng-Prusoff equation (**Equation S4**), modified for allosteric assembly inhibition induced by orthosteric inhibitors. The IC_50_ and K_i_ values in the presence of substrate revealed an increasing inhibitory potency trend from C5 to C13 and C15. In absence of the substrate, only C15 yielded an exact IC_50_ value.

**Table 1.**
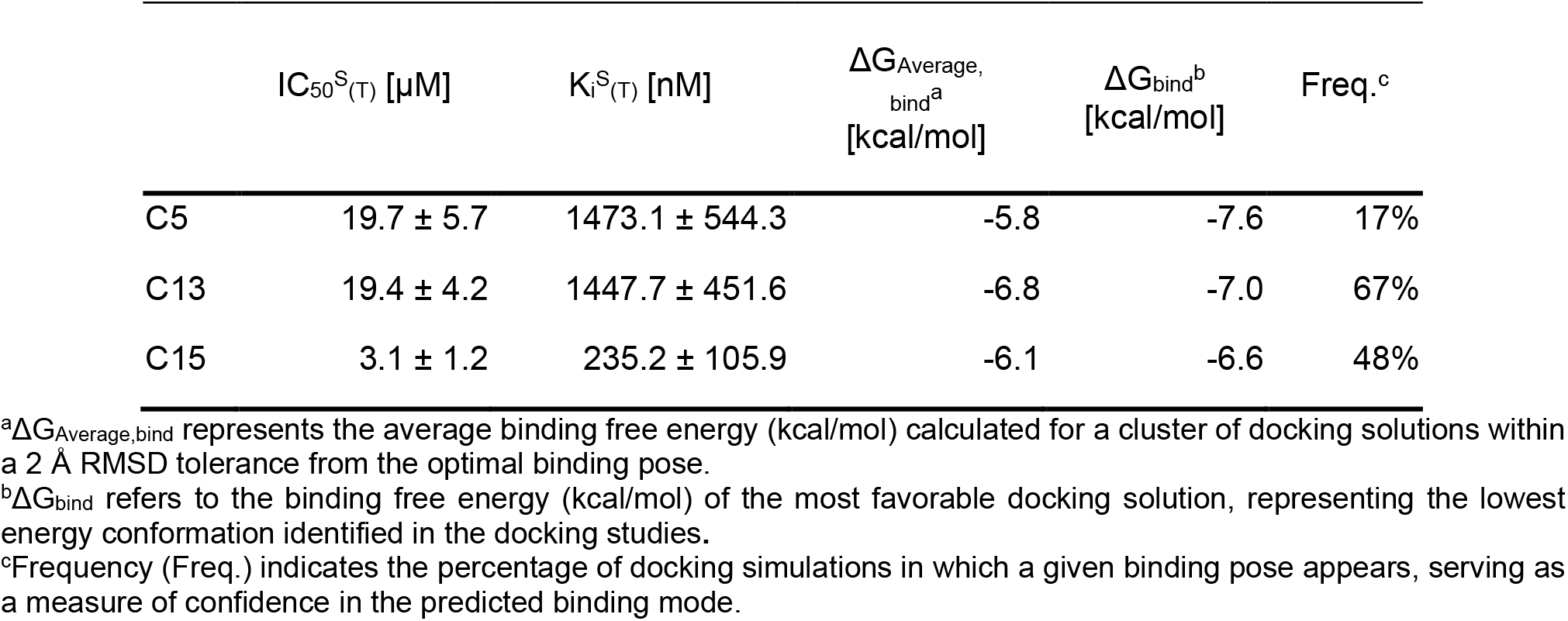
Quantitative analysis of inhibitor potency and binding energies for GNE oligomer disruption. The table summarizes experimentally and computationally derived parameters for the inhibition of GNE tetramers (T) by small molecules C5, C13, and C15. IC_50_ and K_i_ were determined by mass photometry in the presence of substrate (IC_50_^S^_(T)_, K_i_^S^_(T)_), using 100 µM UDP-GlcNAc. Binding energy values (ΔG_Average,bind_, ΔG_bind_) were obtained from molecular docking studies and reflect the predicted interactions between each inhibitor and the GNE binding site. Superscripts S denotes conditions with substrate. Errors represent the standard deviation from three independent experiments, each performed in technical triplicate.

In addition to determining IC_50_ values, we sought to adapt the principles of Schild analysis to estimate the inhibitor affinity (K_B_) for GNE and to gain deeper insight into the underlying inhibitory mechanism. As a representative case, we focused on the most potent inhibitor, C15. For this purpose, we performed substrate titrations at a fixed GNE concentration (800 nM) in the absence or presence of increasing C15 concentrations. The resulting dose-response curves (**Figure 4A**), fitted using **Equation S3** (assuming EC_50_ = IC_50_), showed a progressive rightward shift in EC_50_ with increasing inhibitor concentration, accompanied by a reduction in maximal response amplitude. To quantify the shifts, EC_50_ values obtained in the presence of C15 (EC_50,Inh_) were normalized to the control (EC_50_ in absence of inhibitor), and transformed into concentration ratios (CR), from which CR-1 values were plotted against inhibitor concentration in a double-logarithmic Schild plot (**Figure 4B**). The resulting curve revealed three distinct regions: a no-effect region at low concentrations (2 nM - 0.4 µM), a linear region (LR, 0.8 - 3.2 µM), and a non-linear saturation region (NLR, >3.2 µM). The NLR was fitted using a non-linear Schild equation (**Equation S5**), adapted from literature sources (see Supporting Information).^23^ The resulting curvature indicated a cooperative inhibition mechanism. In contrast, the LR was analyzed independently (**Figure S4A**) using a standard linear Schild equation (**Equation S6**). The observed slope (m = 1.5) deviated from the classical value of 1 expected for purely competitive inhibition, further supporting the presence of cooperative effects. Due to this deviation and the complexity of the inhibition profile, a precise dissociation constant for the inhibitor (K_B_) could not be obtained. Instead, an apparent affinity (K_B,app_ ≈ 1.0 µM) was estimated from the x-intercept of the linear fit (**Figure S4A**).

**Figure 4.**
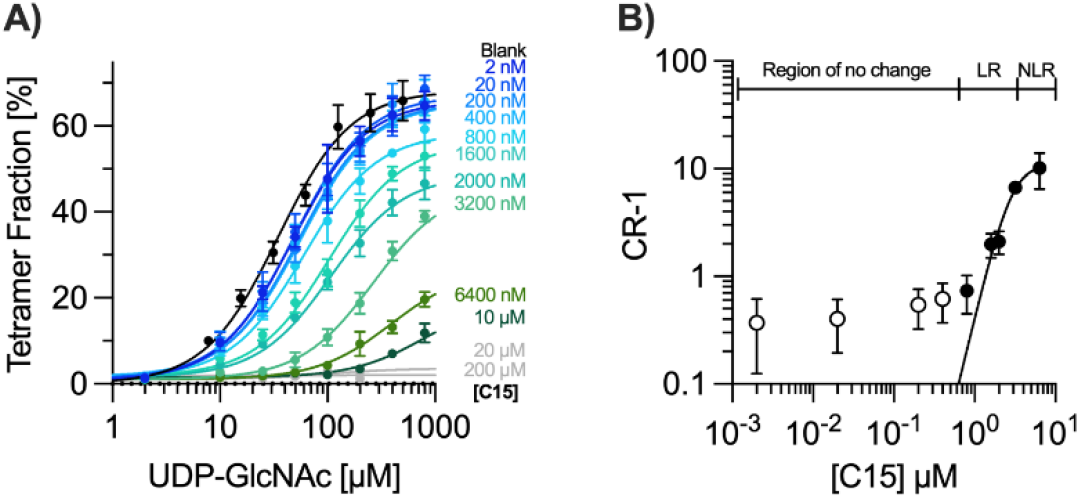
Schild analysis and Schild-plot for GNE tetramer inhibition by C15. A) Mass photometry analysis of GNE tetramer assembly as a function of substrate concentration (UDP-GlcNAc) in the presence of increasing concentrations of inhibitor C15 (indicated by blue to green curves). Curves with no change are colored in grey. Tetramer fractions are shown as percentages of the total protein concentration. Data were fitted using **Equation S3** with a Hill coefficient n = 1.3, as determined in **Figure 2G**. B) Double-logarithmic Schild plot illustrating the inhibition of GNE tetramer assembly by C15. CR-1 values were calculated as the ratio of EC_50_ values in the presence *versus* absence of inhibitor, minus one, based on the substrate titration curves in panel A). Data points (filled circles) were fitted using a modified non-linear Schild-equation (**Equation S5**). The plot is divided into three regions, indicated at the top: a low concentration “region of no change”, a linear region (LR) and a non-linear region (NLR) at high C15 concentrations. All measurements were performed at a fixed GNE concentration of 800 nM. Error bars represent the standard deviation (A) or the standard error of the mean (SEM, B) from six replicates.

To complement this analysis and gain mechanistic insight beyond the linear range, we applied the Hill-modified Operational Model of Allosterically Modulated Agonism (OMAM) (**Equation S7**)^24^ to globally fit the full substrate response curves. Key parameters, including maximal response (E_max_ = 68.3%), substrate affinity (K_S_ = 34.6 µM), and the Hill-coefficient (n = 1.3) were fixed based on prior analysis (**Figure 2G**). From the OMAM fit (Figure S4B), we extracted operational efficacy values: **τ**_S_ = 18.2 for the substrate and **τ**_I_ = 0.02 for the inhibitor, reflecting the strong tetramer-inducing capacity of UDP-GlcNAc and the negligible intrinsic efficacy of C15. Additionally, the model yielded a binding cooperativity factor (α = 0.04), indicating strong negative cooperativity between C15 and substrate binding, consistent with allosteric modulation of the protein. The operational cooperativity factor (β ≈ 0) suggests that C15 nearly abolishes the functional effect of the substrate upon binding, likely by inducing conformational changes that disrupt tetramer formation. This is consistent with the pronounced reduction in maximal response observed in the titration curves (**Figure 4A**). Notably, the inhibitor affinity (K_B_ = 44.6 nM) estimated from the OMAM global fit differs substantially from the apparent K_B,app_ obtained via Schild analysis of the linear region.

### Molecular modeling studies

To atomistically explain the experimentally observed differences in inhibitor potency among C5, C13 and C15, and to gain deeper insights into their modes of action, we performed docking studies of the inhibitors with GNE monomers. Consistent with C5, both C13 and C15 exhibited preferential binding within the same protein pocket as UDP-GlcNAc (**Figure 5**), supporting the inhibition mechanism proposed in our previous study based on HDX-MS analysis and structural similarities between the inhibitors and the substrate.^7^ For each inhibitor, we identified the lowest-energy binding pose and calculated the average binding energy (ΔG_Average,bind (T)_) for a cluster of poses within a 2 Å Root Mean Square Deviation (RMSD) tolerance from the optimal docking solution. The resulting binding energy values (ΔG_Average,bind_, ΔG_bind_), along with their frequency (Freq.) as a confidence metric, are summarized in **Table 1**. The energy values were obtained with considerable sampling frequencies, representing the percentage of simulation steps in which the inhibitor adopted a given binding pose. These results indicate an increasing inhibitor potency trend from C5 to C15 and C13, deviating slightly from our experimental findings.

**Figure 5.**
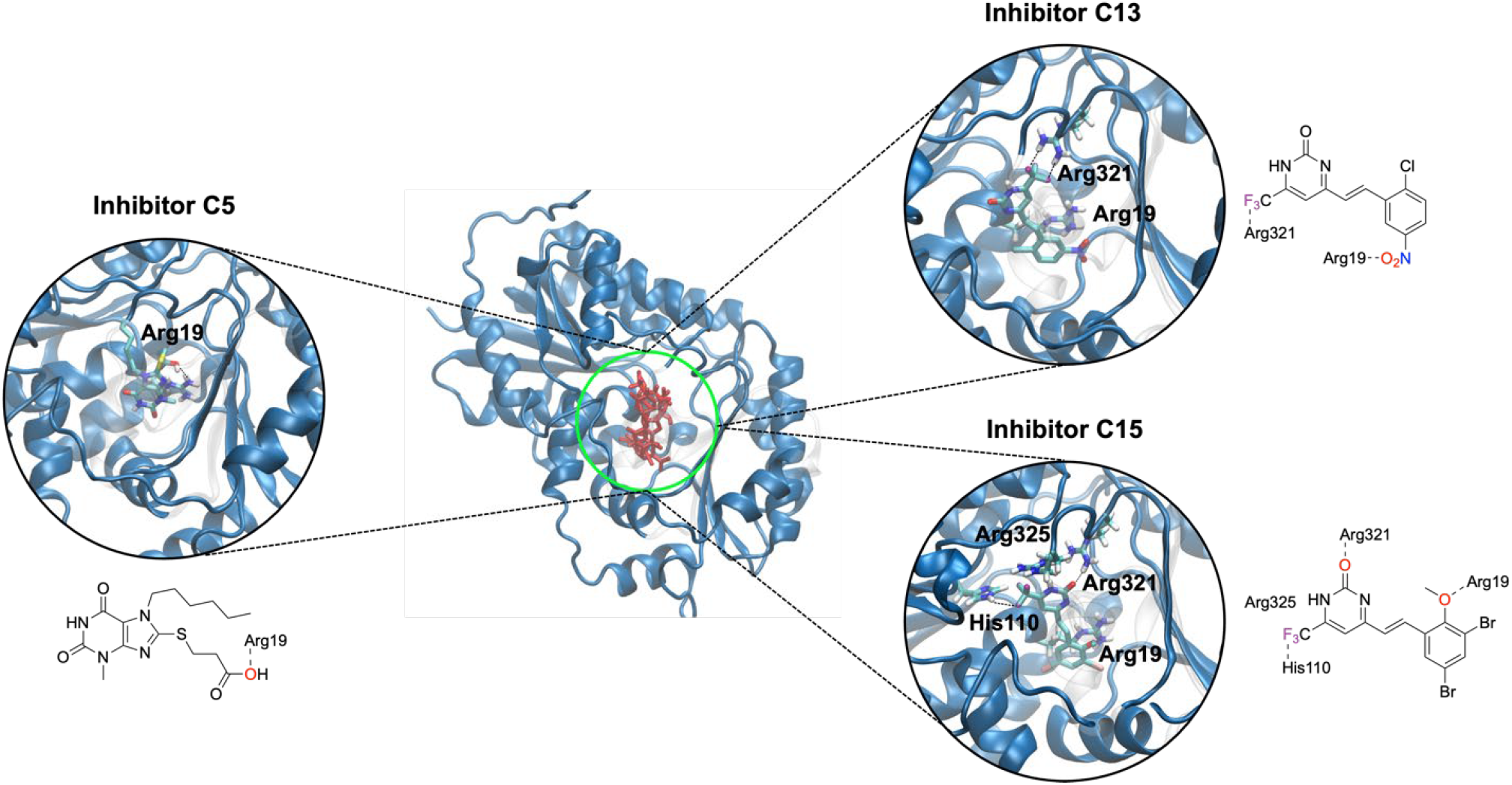
Docking analysis of inhibitor binding to the GNE monomer. The central panel displays the GNE crystal structure (PDB, ID: 4zht),^4^ with an overlay of the lowest-energy, most populated docking poses (red) for all inhibitors. Independent docking results for C5 (left), C13 (top right), and C15 (bottom right) illustrate their respective binding orientation within the enzyme. Key interacting residues (depicted in licorice representation) within the GNE binding pocket are highlighted. The docking results reveal that all inhibitors occupy the substrate binding cleft, overlapping with the UDP-GlcNAc binding site (green circle). The 2D structures of the inhibitors indicate chemical groups involved in interactions with GNE residues.

To further interpret the docking results and gain deeper insight into the inhibition mechanism, we analyzed the key molecular interactions between the inhibitors and residues in GNE’s substrate binding pocket (**Figure 5**). These interactions primarily differ in the number and positioning of hydrogen bonds (H-bonds) formed between enzyme residues and inhibitor molecules, as illustrated in **Figure 5**. The carboxylic oxygen atom of C5 forms an H-bond with Arg19, a key residue in the binding site. Similarly, C13 and C15 engage in H-bonding at Arg19, with the oxygen of the C13 nitro group and the ether group of C15 interacting at this site. Additionally, the trifluoromethyl fluorine atoms of C13 and C15 establish H-bonds with Arg321 and His110, respectively. C15 also forms an H-bond between its 2-pyrimidinone oxygen and Arg321, while the positive charge of Arg325 further stabilizes the interaction.

Notably, the H-bond interactions at Arg19 and Arg321, which are common to all three inhibitors, are also observed in UDP binding to the GNE active site (**Figure 1A**). This suggests a competitive inhibition mechanism, where the inhibitors mimic substrate interactions, thereby blocking UDP-GlcNAc binding and disrupting GNE function.

## DISCUSSION

The enzymatic activity of uridine diphosphate N-acetylglucosamine 2-epimerase (GNE) is inherently linked to its assembly state, with dimerization and tetramerization playing a crucial role in regulating its function.^7^ Due to the dynamic nature of GNE assembly, interferometric scattering microscopy (iSCAM) provides an ideal tool for studying its assembly equilibria and the influence of inhibitors. In this study, we leveraged iSCAM to comprehensively investigate GNE intermolecular stability in the presence or absence of UDP-GlcNAc and to characterize the potency and mechanism of action of three non-carbohydrate-based inhibitors (C5, C13, C15), previously identified in our research.^7^ Computational docking studies further supported these findings by providing atomistic insights into inhibitor binding affinities and potency trends.

Initial GNE titration experiments using iSCAM revealed three distinct molecular assemblies corresponding to monomers (∼50 kDa), dimers (∼90 kDa), and tetramers (∼180 kDa). The absence of trimeric intermediates suggests a stepwise assembly mechanism, transitioning from monomers to dimers and subsequently to tetramers, consistent with previous reports.^4,7^ In the absence of UDP-GlcNAc, GNE predominantly existed as dimers, with strong monomer-monomer (M:M) affinity in the nanomolar range (K_D_^0^_(M:M)_ = 10.9 ± 1.0 nM) and weaker dimer-dimer (D:D) interactions in the micromolar range (K_D_^0^_(D:D)_ = 23.5 ± 4.8 µM). Substrate addition markedly shifted the equilibrium towards tetramer formation, increasing D:D affinity by two orders of magnitude and promoting strong positive cooperativity, as reflected by a Hill coefficient > 1. Quantitative analysis of substrate titration data at a fixed GNE concentration (800 nM) further confirmed moderate substrate binding affinity (K_S_ = 34.6 ± 3.0 µM), and cooperativity, corroborating earlier HDX-MS findings.^7^

To investigate the inhibitory mechanisms of C5, C13, and C15, we previously assessed their effects on GNE dimer and tetramer stability at a single high inhibitor concentration in the presence of UDP-GlcNAc. These semi-quantitative comparisons suggested a potency ranking of C5 < C13 < C15, consistent with analytical size-exclusion chromatography (aSEC) data and enzymatic activity assays.^7^ Subsequent inhibitor titration experiments allowed us to derive IC_50_ values (**Table 1, S1**), confirming this trend. Assuming a mixed competitive and allosteric inhibition mechanism, suggested by HDX-MS data,^7^ molecular docking, and the observed Hill-coefficient, we used a modified Cheng-Prusoff equation (**Equation S4**) to convert IC_50_ values into K_i_ values.

To gain further mechanistic insight, we modeled the concentration-dependent inhibition data for the most potent inhibitor, C15. Modified Schild analysis revealed three regimes: no effect at low concentrations, a linear range with slope m > 1 suggesting cooperativity, and a saturation plateau at high concentrations, indicative of complex binding dynamics. From the linear regime we could determine an apparent inhibitor affinity (K_B,app_) with a value of ∼1.0 µM. However, to more accurately capture the cooperative and allosteric behavior observed across the full dose-response range, we applied the Operational Model of Allosterically Modulated Agonism (OMAM; **Equation S7**) to globally fit the substrate-response curves.^24^

This model yielded efficacy (**τ**) and cooperativity (α, β) parameters: C15 showed minimal intrinsic efficacy (**τ**_I_ = 0.02) and strong negative binding cooperativity (α = 0.04), with almost complete suppression of substrate efficacy (β ≈ 0), consistent with the observed amplitude reduction in the titration curves. The inhibitor affinity derived from the OMAM fit (K_B_ = 44.6 nM) was ∼ 20-fold lower than the K_B,app_ obtained from the linear Schild analysis. This discrepancy underscores not only the limitations of Schild analysis in systems with cooperative interactions but also the fact that both affinity values are model-based estimates. While OMAM provides a mechanistic and quantitative interpretation of such inhibition, definitive determination of inhibitor affinity would require direct structural or biophysical approaches, such as NMR titration or isothermal titration calorimetry (ITC).

Molecular docking supported a competitive binding mechanism, with all inhibitors occupying the UDP-GlcNAc binding site and forming key interactions with the residues Arg19 and Arg321 – both critical for substrate recognition. The number of hydrogen bonds formed correlated with the inhibitor potency, with C5, C13, and C15 forming one, two, and three stabilizing interactions, respectively, consistent with experimental trends.

In summary, this study provides a quantitative and mechanistic understanding of GNE assembly dynamics and its inhibition by non-carbohydrate-based inhibitors (C5, C13, C15). By confirming the competitive nature of inhibition and identifying a cooperative component influencing GNE assembly stability, we expand upon the findings of our previous work.^7^ This work not only advances the mechanistic understanding of GNE regulation but also highlights the utility of iSCAM and OMAM modeling for probing dynamic protein assembly and inhibitor mechanisms at the single-molecule level.

## RESOURCE AVAILABILITY

### Lead contact

For further information or requests for resources and reagents, please contact Prof. Dr. Daniel Lauster.

## ACKNOWLEDGMENTS

This project was conducted with the support of research infrastructure provided by the Research Building SupraFAB, funded by the Federal Government (BMBF) and the State of Berlin. DCL gratefully acknowledges financial support from the Federal Ministry of Education and Research through the project MucPep (FKZ: 13XP511). The authors also acknowledge funding from the Collaborative Research Center “Dynamic Hydrogels at Biological Interfaces” (CRC 1449), supported by the Deutsche Forschungsgemeinschaft (DFG, German Research Foundation) – Project ID 431232613 – SFB 1449 (sub-projects C01, C04, C05N and IRTG). NB is supported by the Add-on Fellowship provided by the Joachim Herz Foundation. Additionally, we thank Dr. Rumiana Dimova (MPIKG Berlin) and her team for kindly providing glycinin for our experiments. S.D.L thanks Freie Universität Berlin for being appointed in the frame of the Global Faculty Program and acknowledges ANPCyT in Argentina for PICT2021GRF-T1-00403 and PIP Conicet 11220210100256CO.

## AUTHOR CONTRIBUTIONS

Conception and design of the study: D.C.L. Acquisition, analysis, and interpretation of the data: N.B., D.C.L., S.D.L. The manuscript was written through contributions of all authors (N.B., D.C.L., C.P.R.H., S.D.L., J.L.). All authors have given approval to the final version of the manuscript.

## DECLARATION OF GENERATIVE AI AND AI-ASSISTED TECHNOLOGIES

During the preparation of this work, the authors did not use any AI tools.

## SUPPLEMENTAL INFORMATION

Equation S1-S7, Figure S1-S4, Derivations S1-S3

## KEY RESOURCES TABLE

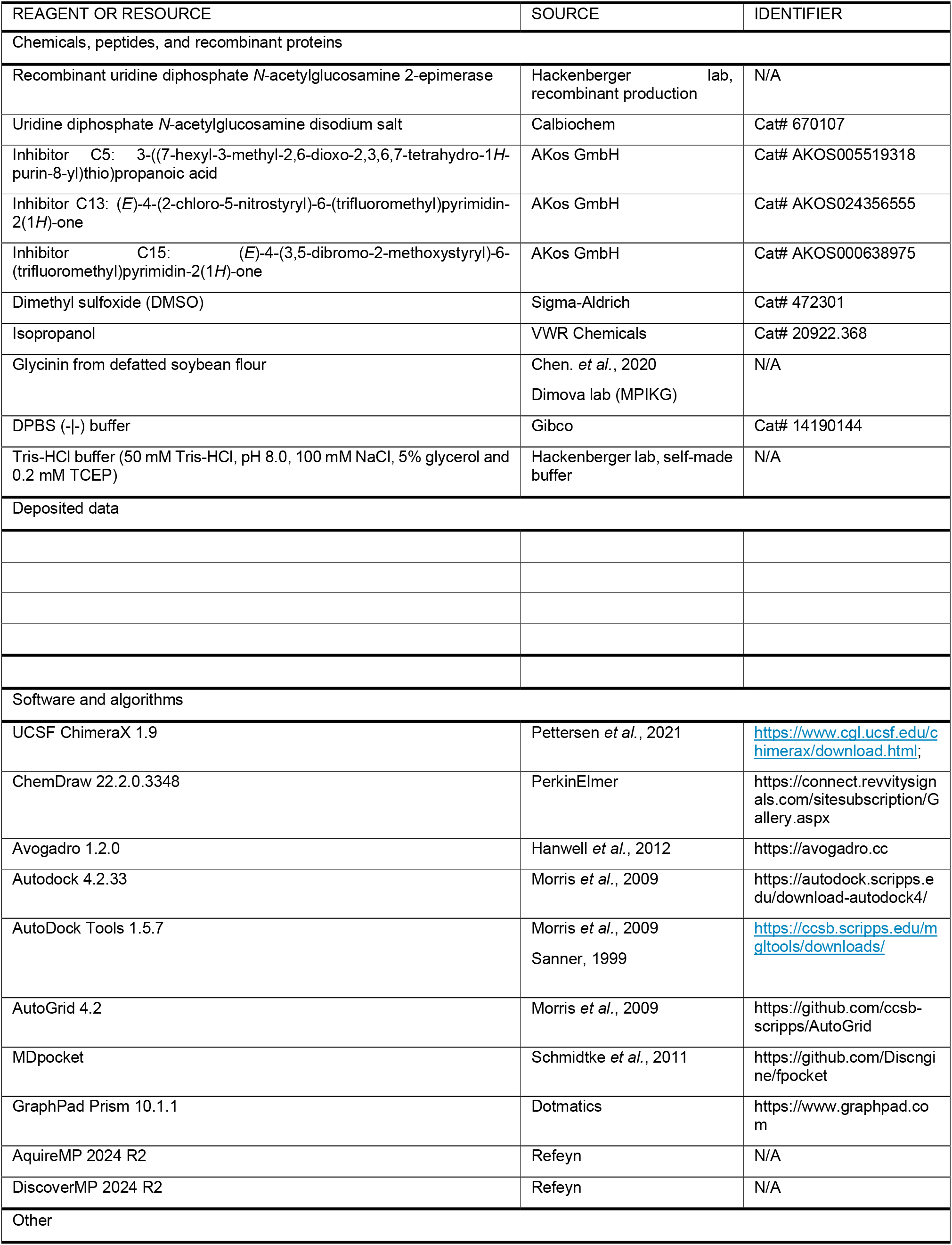
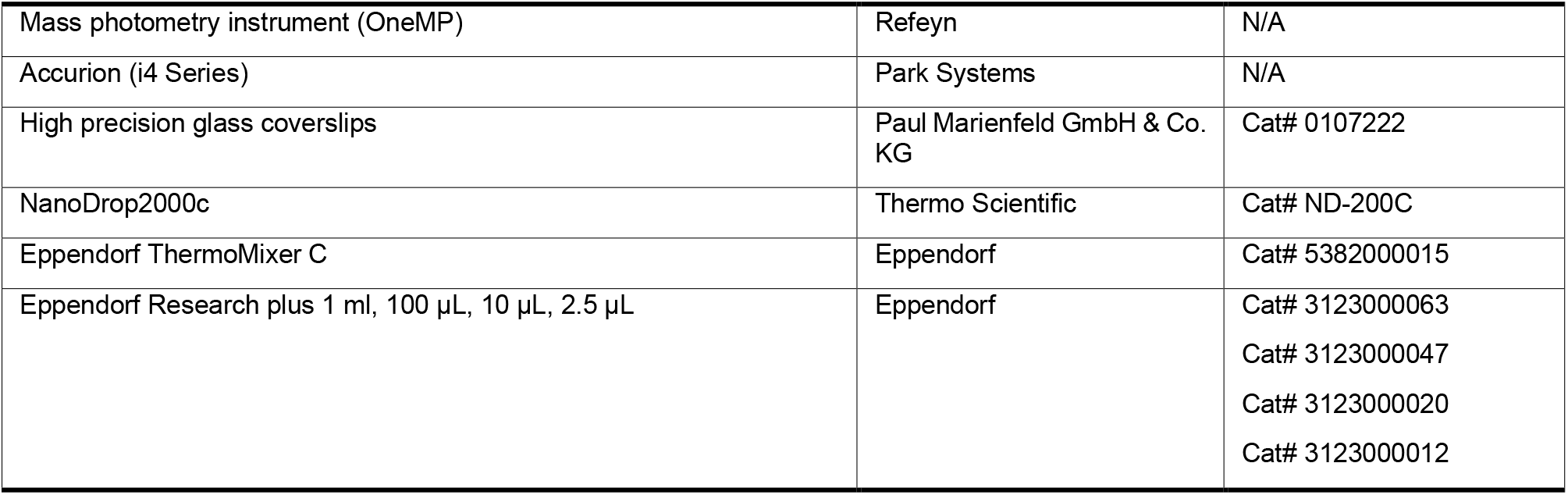

## METHOD DETAILS

### GNE expression and purification

The expression and purification of human UDP-GlcNAc 2-epimerase (GNE) were carried out following the protocol established by the Chen group.^4^ The GNE protein used in this work was expressed as a GNE with an *N*-terminal His_6_-tag in *Escherichia coli* BL21 (DE3) cells. Details on protein production and purification have been previously described in Gorenflos López *et al*..^7^

### UDP-GlcNAc, C5, C13, C15 and GNE sample preparation

GNE sample aliquots were stored frozen at -80 °C in Tris-HCl buffer (50 mM Tris-HCl, pH 8.0, 100 mM NaCl, 5% (v/v) glycerol, and 0.2 mM TCEP). Prior to use, samples were centrifuged (10 minutes, 30,000 x g, 4 °C), and the protein concentration was determined using a NanoDrop2000c (Thermo Scientific) at 280 nm. For concentration determination, the experimentally determined molar extinction coefficient (ε = 19,535 M/cm) and the molecular mass of the His_6_-tagged monomer (46.4 kDa) were used.^6,7^ UDP-GlcNAc (Calbiochem) was prepared as a 10 mM stock solution in DPBS (−|-) buffer (Gibco). Inhibitors C5, C13 and C15 were dissolved in DMSO as 10 mM stock solutions. All stock solutions were stored at - 20 °C until use.

### Mass Photometry (MP) experiments

All mass photometry (MP) measurements were conducted using a OneMP (Refeyn) instrument equipped with a 525 nm green laser. To minimize background noise, the instrument was placed on an active vibration isolation system (i4 Series, Accurion). High-precision glass coverslips (Paul Marienfeld GmbH & Co. KG) were thoroughly cleaned before use to ensure optimal measurement conditions. Prior to MP measurements, samples were incubated in phosphate buffer (DPBS(−|-), Gibco) and low-binding tubes for 30 minutes at 20 °C to allow binding equilibrium to be reached. Immediately before measurement, a 50 nM working solution was prepared. Then, 5 µL of the working solution was added to a 15 µL DPBS droplet on the glass coverslip, followed by mixing upon pipetting to ensure proper distribution before measurements began immediately. This approach provided high reproducibility in our results. For the measurements a regular field of view was used. Data acquisition was performed over a 60-seconds detection period with manual focus adjustment. Data were recorded using the software AquireMP and referenced against a glycinin mass standard in DPBS (−|-). The glycinin standard, derived from defatted soybean flour, forms characteristic trimers (160 kDa), hexamers (320 kDa), nonamers (480 kDa), and dodecamers (640 kDa). Purified and lyophilized glycinin was kindly provided by the group of Dr. Rumiana Dimova.^25,26^ Data analysis and graphical visualization were performed using DiscoverMP and GraphPad Prism 10. All experiments were conducted in at least three independent replicates, each performed as technical triplicates.

### Affinity measurements of GNE subunits in the presence or absence of UDP-GlcNAc

To determine the intermolecular protein affinities of GNE subunits, their assembly distribution was assessed using iSCAM experiments. In the absence of substrate, serial GNE dilutions (50 µM to 50 nM) were performed to evaluate GNE subunit interactions. In the presence of UDP-GlcNAc, intermolecular affinity changes were assessed through two titration approaches: First, different concentrations of UDP-GlcNAc (500 - 7.81 µM) were titrated against a fixed concentration of GNE (0.8 µM). Second, different concentrations of GNE (0.2 - 10 µM) were titrated against a fixed concentration of UDP-GlcNAc (100 µM).

K_D_ values were obtained from binding curves (**Figure S1**), fitted using a Hill-based logistic model (**Equation S1**).^13^ The following dissociation constants were determined: K_D_^0^_(M:M)_ and K_D0(D:D)_ in the absence of UDP-GlcNAc with a y-axis intercept of zero, K_D_^S^_(D:D)_ in the presence of UDP-GlcNAc, also with a y-axis intercept of zero, and K_D_^S^_(M:M)_ with a y-axis intercept defined by the dimer fraction ([GNE_D_]/[GNE_total_]) at 800 nM GNE in the absence of UDP-GlcNAc (**Figure 2C**).

### GNE assembly inhibition experiments

The effects of inhibitors (C5, C13 and C15) on GNE assembly were evaluated using iSCAM. Two-fold serial dilutions of the inhibitors (500 - 7.81 µM) were titrated against a constant GNE concentration (800 nM), either in the presence or absence of UDP-GlcNAc (100 µM). The GNE and substrate concentration were selected based on titration experiments ensuring binding saturation and the establishment of an assembly equilibrium. Prior to measurements, samples were incubated for 30 minutes at 20 °C to allow equilibrium to be reached. IC_50_ values were determined from protein fraction *vs*. inhibitor concentration plots and fitted using a Hill-based logistic function (**Equation S3**). Subsequently, K_i_ values for GNE inhibitors were derived from a Cheng-Prusoff equation accounting for allostery (**Equation S4**), detailed in the Supporting Information.

### GNE-inhibitor Schild experiments

To assess the binding affinity of inhibitors to GNE, modified Schild experiments were performed using iSCAM. Serial dilutions of UDP-GlcNAc (800 µM - 100 nM) were titrated against a constant GNE concentration (800 nM) in the presence of varying concentrations of inhibitor C15 (200 µM - 2 nM). Enzyme and substrate concentrations were chosen based on prior titration experiments (**Figure 2C, G**) to ensure a stable oligomeric equilibrium. Inhibitor concentrations were selected around the IC_50_ to enable accurate response characterization. Samples were incubated for 30 minutes at 20 °C to ensure equilibration before measurement. EC_50_ values were obtained from substrate titration curves (tetramer fraction vs. UDP-GlcNAc concentration), fitted using a Hill-based function (**Equation S3**). The resulting EC_50_ values in the presence of inhibitor (EC_50,Inh_) were converted into concentration ratios (CR-1) and plotted against the corresponding inhibitor concentrations to generate Schild plots. Data points were then fitted using modified non-linear (**Equation S5**) and linear (**Equation S6**) Schild equations, as described in the Supporting Information. Further cooperative factors were obtained from fitting the substrate concentration dependent response curves with an OMAM model (**Equation S7**).^24^

### Docking studies

Molecular docking studies were performed using Autodock 4.2.33,^27^ by using the crystal structure of GNE (PDB ID: 4zht).^4^ Ligand coordinates were generated using Avogadro 1.2.0.^28^ Both proteins and ligands were preprocessed with AutoDock Tools (ADT) package to merge nonpolar hydrogens, calculate Gasteiger charges, and select the rotatable side-chain bonds.^27^ For docking evaluation, grid boxes were set with a spacing of 0.375 Å and dimensions of 60 × 60 × 60 points, centered on a binding pocket identified using the MDpocket program.^29^ Grid maps were subsequently generated using AutoGrid 4.2, which is included in the AutoDock 4.2 distribution.^27^ All docking simulations were conducted using default AutoDock parameters.^27^

