## Supporting Information for "Quantitative analysis of inhibitor-induced assembly disruption in human UDP-GlcNAc 2-epimerase using mass photometry"

$$f([GNE]) = M_1 + (M_2 - M_1)\left(\frac{[GNE]^n}{([GNE]^n + K_D{}^n)}\right)$$

**Equation S1. Hill-based logistic fitting function for determination of dissociation constants ($K_D$).** This equation describes the protein fraction f([GNE])) as a function of GNE protein concentration ([GNE]), where f([GNE]) represents the fraction of a specific GNE assembly state (X), normalized to the total GNE concentration ($[GNE_X]/[GNE_{total}]$), where X denotes monomer (M), dimer (D), or tetramer (T). $M_1$ and $M_2$ define the lower and upper plateau regions of the protein fraction. $K_D$ represents the dissociation constant, quantifying the equilibrium between free and assembled protein, and n = 1.3 is the Hill coefficient, determined in **Figure 2G** (main article), indicating the degree of cooperativity in the assembly process. The equation is based on the law of mass action and was modified from Fineberg *et al.*.[1]

$$f([S]) = \frac{E_{max} \cdot [S]^n}{K_S{}^n \cdot [S]^n}$$

**Equation S2. Hill function for the determination of substrate affinity to the enzyme and maximal response in substrate titration experiments in absence of inhibitor.** This equation models the protein fraction f([S]) as a function of the substrate concentration [S]. $E_{max}$ (68.3%) represents the maximal possible response (in our case maximal percentage of tetramer fraction under defined conditions), $K_S$ the substrate affinity to the enzyme and n = 1.3 the Hill-coefficient, determined in **Figure 2G** (main article).

$$f([I]) = M_1 + (M_2 - M_1)\frac{1}{1 + \left(\frac{IC_{50}}{[I]}\right)^n}$$

**Equation S3. Hill-based fitting function for $IC_{50}$ determination.** The equation models the protein fraction f([I]) of individual protein subunits (M = monomer, D = dimer, T = tetramer) as a function of the inhibitor concentration ([I]). $M_1$ and $M_2$ define the minimal and maximal plateau regions of the fraction of assembled protein. $IC_{50}$ represents the half-maximal inhibitory concentration and (n = 1.3) the Hill-coefficient, determined in **Figure 2G** (main article). This function is derived from the rate law of mass action, as detailed in **Derivation S1**.

$$K_i = \left(\frac{IC_{50}}{1 + \left(\frac{[S]}{K_{0.5}}\right)^n}\right) \cdot \alpha$$

**Equation S4. Modified Cheng-Prusoff equation for the $IC_{50}$ conversion into $K_i$ for allosteric protein assembly inhibition.** Modified equation for converting $IC_{50}$ values into assay-independent inhibition constants ($K_i$) for competitive and allosteric protein assembly inhibition. $IC_{50}$ denotes the inhibitor concentration required for half-maximal inhibition, determined from titration experiments (**Table 1, S1**). $K_{0.5}$ represents the half-maximal assembly saturation ($K_{0.5,(M:M)}$ = 10.9 nM, $K_{0.5,(D:D)}$ = 23.5 μM), derived from GNE dilution assays. [S] is the substrate concentration used (100 μM), n is the Hill coefficient (n = 1.3), determined in **Figure 2G** (main article), and

$\alpha$ is a correction factor ($\alpha_D$ = 0.3, $\alpha_T$ = 0.5) accounting for the oligomeric nature of the protein mixture. The derivation and modification of the equation are provided in **Derivation S2**.

$$f([I]) = \left(\frac{1 - \lambda}{\lambda}\right)\left(\frac{[I]^n}{\left(\frac{K_B}{\lambda}\right) + [I]^n}\right)$$

**Equation S5. Modified non-linear Schild fitting function.**
This equation, adapted as a modified version from Lane *et al.*,[2] models the non-linear region of the Schild plot. f([I]) represents the competitive concentration ratio parameter (CR-1), defined as the ratio of the $EC_{50}$ in the presence of inhibitor ($EC_{50,Inh}$) to the $EC_{50}$ in its absence, minus one. It describes the influence of an allosteric, orthosteric inhibitor on protein assembly in the presence of the substrate (UDP-GlcNAc) across varying inhibitor concentrations. The term $\lambda$ denotes the composite cooperativity factor, consisting the substrate-inhibitor cooperativity and the effect on the signaling efficacy.[2] $K_B$ represents the apparent dissociation constant of the inhibitor and n the Hill-coefficient (n = 1.3) determined in **Figure 2G** (main article). The full derivation is provided in **Derivation S3.**

$$\log(CR\text{-}1) = f([I]) = m \log([I]) - m \log(K_B)$$

**Equation S6. Linear Schild fitting function.**
This equation, adapted in modified form from Lane *et al.*,[2] describes the linear region of the Schild plot. f([I]) represents the logarithmic form of the competitive concentration ratio parameter, log(CR-1), where CR is defined as the ratio of $EC_{50}$ in the presence ($EC_{50,Inh}$) to that in the absence of inhibitor. The function characterizes the effect of an inhibitor on protein assembly in the presence of the substrate (UDP-GlcNAc) across varying inhibitor concentrations ([I]). m denotes the slope of the linear fit (typically $m > 1$ in cooperative systems) and $K_B$ is the apparent dissociation constant of the antagonist. A detailed derivation is provided in **Derivation S3**.

$$f([S]) = E = \frac{E_{max} \cdot (\tau_S \cdot [S]^n \cdot \left(K_B^n + \alpha \cdot \beta \cdot [I]^n\right) + \tau_I \cdot [I]^n \cdot K_S^n)}{[S]^n \cdot K_B^n + K_S^n \cdot K_B^n + [I]^n \cdot K_S^n + \alpha \cdot [S]^n \cdot [I]^n + \tau_S \cdot [S]^n \cdot \left(K_B^n + \alpha \cdot \beta \cdot [I]^n\right) + \tau_I \cdot [I]^n \cdot K_S^n}$$

**Equation S7. Hill modified operational model of allosterically modulated agonism (OMAM) for the modelling of Schild experiments.**
This equation describes the response f([S]) = E as a function of the orthosteric substrate concentration ([S]) in presence of varying allosteric inhibitor concentrations ([I]). Cooperativity is described by the factors $\alpha$ (binding cooperativity), $\beta$ (operational cooperativity) and operational efficacies by $\tau_S$ and $\tau_I$ for the substrate and the inhibitor respectively. $E_{max}$ denotes the maximal possible response (maximal tetramer fraction, 68.3%) determined in **Figure 2G** (main article). Dissociation constants are named $K_S$ (34.6 µM) for the substrate, determined in **Figure 2G** (main article), using **Equation S2** and $K_B$ for the inhibitor. n is the Hill-coefficient (n = 1.3) determined in **Figure 2G** (main article), introduced for a better fit of the concentration-response curves. The equation was adapted in modified form from Jakubík *et al.*.[3]

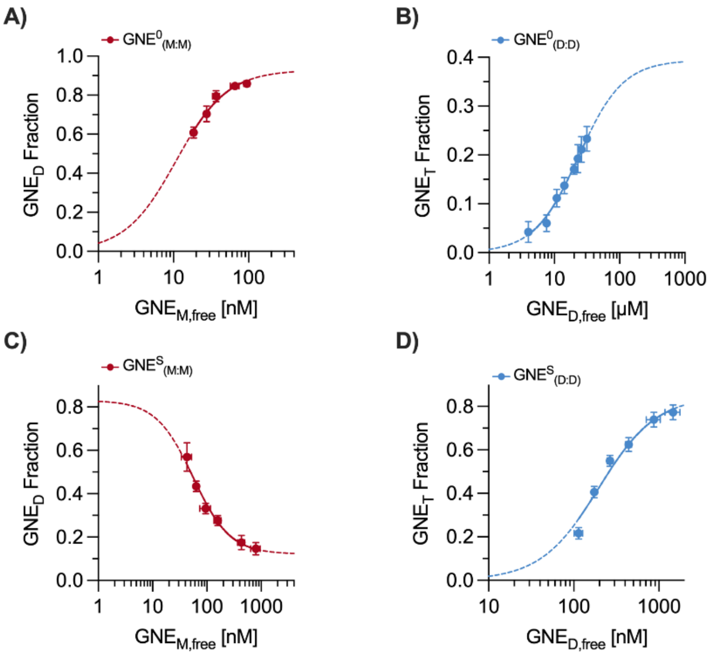

**Figure S1. Mass photometry analysis of GNE dimer and tetramer fractions at increasing monomer concentration.**

Binding curves illustrating the dissociation constants ($K_D$) for GNE subunit interactions (monomer-monomer (M:M, red), dimer-dimer (D:D, blue)) in presence (S) or absence (0) of UDP-GlcNAc. A) Binding curve for monomer-monomer interactions in the absence of UDP-GlcNAc, yielding $K_D{}^0{}_{(M:M)}$ for dimer formation. B) Binding curve for dimer-dimer interactions in the absence of UDP-GlcNAc, yielding $K_D{}^0{}_{(D:D)}$ for tetramer formation. C) Binding curve for monomer-monomer interactions in the presence of UDP-GlcNAc (S), yielding $K_D{}^S{}_{(M:M)}$. D) Binding curve for dimer-dimer interactions in the presence of UDP-GlcNAc (S), yielding $K_D{}^S{}_{(D:D)}$. The GNE fraction represents the proportion of dimer or tetramer populations relative to the total protein concentration in the analyzed solution. $GNE_{M,free}$ and $GNE_{D,free}$ refer to the concentrations of free monomers and dimers, respectively. Error bars indicate the standard deviation from three independent experiments, each performed as technical triplicates. Due to the logarithmic scale**,** error bars in the x-direction may not always be visible**.** Binding curves were fitted using **Equation S1**, with a Hill-coefficient of n = 1.3 determined in **Figure 2G** (main article).

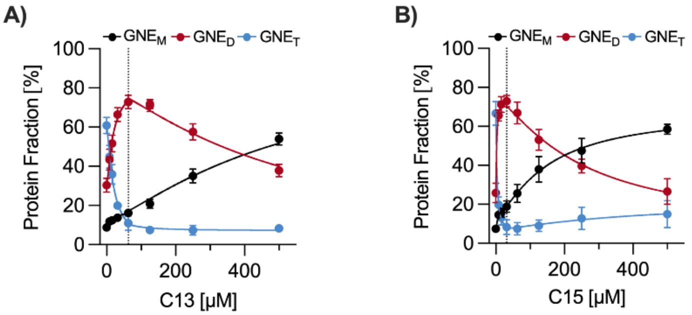

**Figure S2. Concentration-dependent effects of C13 and C15 on GNE assembly at constant UDP-GlcNAc levels.**

Effect of inhibitor concentration on GNE assembly, shown as the protein fraction of each subunit: monomer (M, black), dimer (D, red), tetramer (T, blue) expressed as a percentage of the total protein concentration. Impact on GNE assembly is determined for A) C13, and B) C15. The dotted grey lines separate low and high concentration regimes, which were analyzed individually (see **Figure 3** in the main article). Each concentration regime was fitted independently using **Equation S3**, with a Hill-coefficient of n = 1.3 determined in **Figure 2G** (main article). Error bars indicate the standard deviation from three independent experiments, each performed as technical triplicates.

**Table S1. Inhibitory potency of C5, C13, and C15 against GNE dimers in the presence and absence of UDP-GlcNAc.**

Experimentally and computationally derived potency values ($IC_{50}$ and $K_i$) for GNE dimer (D) inhibition by C5, C13, and C15. Measurements were performed using mass photometry in the presence ($IC_{50}{}^{S}{}_{(D)}$, $K_i{}^{S}{}_{(D)}$) or absence ($IC_{50}{}^{0}{}_{(D)}$) of 100 µM UDP-GlcNAc. S and 0 denote conditions with and without substrate, respectively. Reported errors represent the standard deviation from three independent experiments, each conducted in technical triplicate.

| | $IC_{50}{}^{S}{}_{(D)}$ [µM] | $K_i{}^{S}{}_{(D)}$ [nM] | $IC_{50}{}^{0}{}_{(D)}$ [µM] |
|---|---|---|---|
| C5 | - | - | - |
| C13 | 500.2 ± 72.8 | 1.4 ± 0.3 | >1000 |
| C15 | 245.4 ± 60.3 | 0.7 ± 0.2 | 23.6 ± 7.2 |

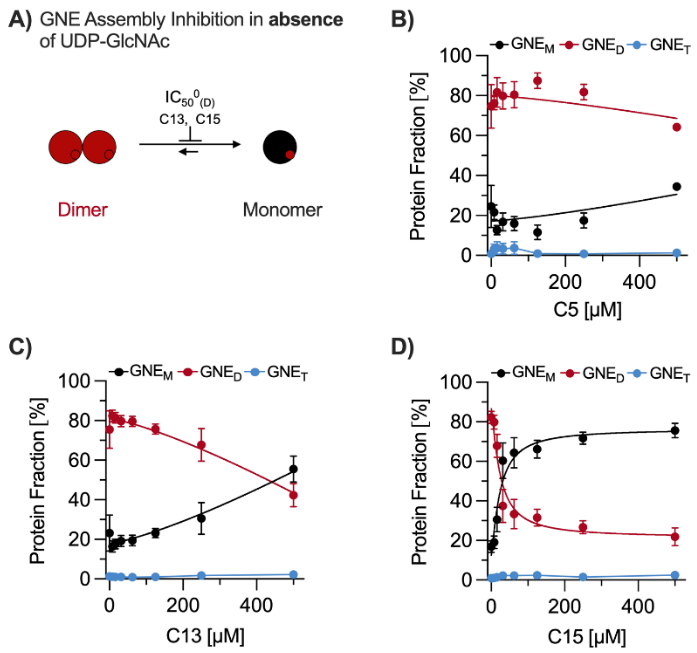

**Figure S3. Concentration-dependent GNE assembly inhibition in the absence of UDP-GlcNAc with increasing concentrations of inhibitors C5, C13 or C15.**

A) Schematic representation of the inhibitor effect (small red spheres) on the monomer-dimer equilibrium in absence of UDP-GlcNAc. Arrows indicate a shift towards the monomer population upon inhibitor binding. $IC_{50}$ values were determined from the following titration experiments. Concentration-dependent effects of inhibitors B) C5, C) C13, D) C15 on GNE subunit distribution, represented as the protein fraction of each subunit: monomer (M, black), dimer (D, red), and tetramer (T, blue) as a percentage of the total protein concentration. Error bars indicate the standard deviation from two independent experiments, each performed as technical triplicate. Curves were fitted using **Equation S3**, with a Hill-coefficient of n = 1.3 determined in **Figure 2G** (main article). Data points of the tetramer populations, that were not possible to fit, due to a too low fractions, were connected *via* strait lines.

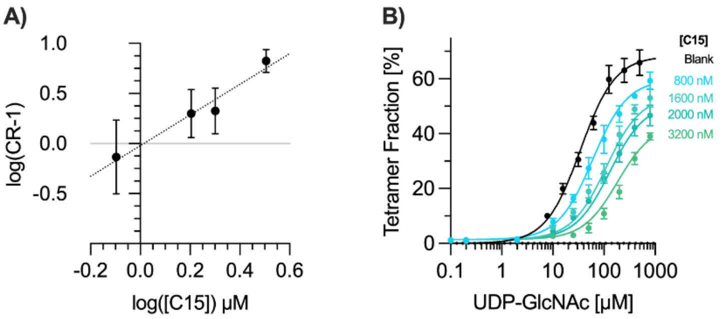

**Figure S4. Quantitative analysis of cooperative inhibition of GNE tetramer formation by C15**

A) Double-logarithmic Schild plot of the linear region (LR) illustrating C15-mediated inhibition of GNE tetramer assembly. CR-1 values were calculated from $EC_{50}$ shifts observed in substrate titration curves (**Figure 4A**), and the LR was fitted using a linear Schild equation (**Equation S6**). The x-intercept corresponds to the apparent $\log(K_{B,app})$. Error bars represent the SEM from six replicates. B) Substrate-dependent GNE tetramer formation analyzed by mass photometry and fitted using a Hill-modified Operational Model of Allosterically Modulated Agonism (OMAM, **Equation S7**) to extract efficacy ($\tau_S$, $\tau_I$) and cooperative parameters (α, β).[3] A fixed maximal response (Emax = 68.3%), substrate affinity ($K_A$ = 34.6 μM), and Hill coefficient (n = 1.3), determined in **Figure 2G** (main article), was used. Error bars represent the standard deviation of six replicates.

### DERIVATIONS

#### Derivation S1. Hill-based inhibition function

The inhibition of protein assembly (A) can be described by a simplified binding equilibrium. In the case of GNE, A denotes either the dimeric or tetrameric form of the enzyme. The inhibitor is represented by I, and m indicates the number of inhibitor molecules that bind to the assembly.

$$A + m\,I \leftrightharpoons AI_m$$

Applying the law of mass action at thermodynamic equilibrium yields the following expression, where the ratio of the inhibitor-bound protein assembly ($[AI_m]$) to the unbound components [A] and [I]) defines the dissociation constant:

$$K_D = \frac{[A]\,[I]^m}{[AI_m]} \qquad (1)$$

To account for cooperative binding, arising from multiple inhibitor binding sites within the assembly, the Hill coefficient (n) is introduced:

$$K_D = \frac{[A]\,[I]^n}{[AI_m]} \qquad (2)$$

Rearranging for the concentration of the inhibitor-bound assembly gives:

$$[AI_m] = \frac{[A]\,[I]^n}{K_D} \qquad (3)$$

Assuming that the total concentration of the assembly remains constant throughout the experiment, the following mass balance equation applies:

$$[A]_{total} = [A] + [AI_m] \qquad (4)$$

Substituting equation (3) into (4):

$$[A]_{total} = [A] + \frac{[A]\,[I]^n}{K_D} \qquad (5)$$

Factoring out [A]:

$$[A]_{total} = [A]\left(1 + \frac{[I]^n}{K_D}\right) \qquad (6)$$

The fraction of inhibitor-bound protein assembly relative to the total is then given by:

$$\frac{[A_m]}{[A]_{total}} = \frac{\frac{[I]^n}{K_D}}{1 + \frac{[I]^n}{K_D}} \qquad (7)$$

Simplifying:

$$\frac{[AI_m]}{[A]_{total}} = \frac{[I]^n}{K_D + [I]^n} \qquad (8)$$

To define the half-maximal inhibitory concentration ($IC_{50}$), we introduce the condition, where 50 % of the protein assembly is inhibited. At this inhibitor concentration, $[I] = IC_{50}$.

Considering cooperative binding, the relationship between $IC_{50}$ and $K_D$ is given by $IC_{50}^n = K_D$. Substituting $K_D$ in equation (8) and dividing numerator and denominator by $[I]^n$ yields the inhibitory form of the Hill equation:

$$\frac{[AI_m]}{[A]_{total}} = \frac{1}{1 + \left(\frac{IC_{50}}{[I]}\right)^n} \qquad (9)$$

To allow for sigmoidal fitting of experimental data, two scaling parameters, $M_1$ and $M_2$, are introduced to define the minimum and maximum plateaus of the inhibition curve, respectively. The resulting equation describes the fraction of inhibited protein assemblies as a function of inhibitor concentration, allowing for determination of $IC_{50}$:

$$f([I]) = M_1 + (M_2 - M_1)\,\frac{1}{1 + \left(\frac{IC_{50}}{[I]}\right)^n} \qquad (10)$$

**Derivation S2. Modification of the Cheng-Prusoff equation considering competitivity and cooperativity.**

The Cheng-Prusoff equation is used to translate $IC_{50}$ values, obtained in enzyme activity inhibition studies, into assay-independent, comparable $K_i$ values. The regular Cheng-Prusoff equation considering competitive inhibition requires the half-maximal inhibitory concentration ($IC_{50}$), the substrate concentration ([S]) and the Michaelis-Menten constant ($K_M$). To apply on assembly inhibition, the $K_M$ can be replaced by the affinity constant $K_D$, derived from substrate titration experiments.

$$K_i = \frac{IC_{50}}{1 + \frac{[S]}{K_M}} = \frac{IC_{50}}{1 + \frac{[S]}{K_D}} \qquad (1)$$

Cooperativity requires the implantation of the Hill-coefficient (n), since the binding behavior is not following the classic Michaelis-Menten model. Further an additional oligomeric correction factor (α) has to be included, considering the ratio of aimed assembly (dimer, tetramer) to the total protein concentration. The affinity measure is included with the half-maximal substrate saturation ($K_{0.5}$).

$$K_i = \frac{IC_{50}}{\left(1 + \frac{[S]}{K_{0.5}}\right)^n} \cdot \alpha \qquad (2)$$

**Derivation S3. Determination of inhibitor affinities ($K_B$) accounting for allostery using adapted Schild plot and function**

To quantify a mixed competitive and allosteric inhibition of GNE assembly in a manner, independent of substrate concentrations, we employed the Schild equation, which is traditionally used to determine antagonist affinity for receptors in presence of an agonist, to derive the binding constant of an antagonist ($K_B$).

The Schild equation uses the concentration ratio parameter (CR-1) to quantify the competitive effect of the antagonist (in our case inhibitor) on the agonist (in our case substrate) potency. The CR-1 parameter is defined as the ratio of the half-maximal effective agonist concentration in the presence ($EC_{50,modulated}$) and absence ($EC_{50,control}$) of the antagonist (in this case, the inhibitors):

$$CR - 1 = \frac{EC_{50,modulated}}{EC_{50,control}} - 1 \qquad (1)$$

For our study, we determined $EC_{50}$ values from Protein Fractions (PF) *vs.* substrate concentration plots, derived from substrate titration experiments in presence or absence of the inhibitor. The PF represent the proportion of monomers, dimers, and tetramers (derived from protein counts) as a percentage of the total protein concentration in solution. The effectivity parameter is defined as the ratio of the $EC_{50,Inh}$ in the presence of the inhibitor (indicated by subscript Inh) to the $EC_{50}$ in absence of the inhibitor, minus one:

$$CR - 1 = \frac{EC_{50,Inh}}{EC_{50}} - 1 \qquad (2)$$

The resulting values were plotted against the inhibitor concentration ([I]).

**Non-linear Schild Equation**

To fit the non-linear part of the Schild-plot, we adopted and modified the Schild equation from literature incorporating cooperative effects and describing agonist-antagonist affinity interactions at receptors.[2]

$$\log(CR - 1) = \log\left(\frac{[B](1 - \alpha\beta)}{\alpha\beta\,[B] + K_B}\right) \qquad (3)$$

The logarithmic transformation used in the Schild plot equation was removed using a base-10 exponentiation for direct fitting. The original equation includes the antagonistic modulator concentration ([B]) and the combined cooperative factor (αβ). In our case, the antagonist modulator (B) is replaced by the inhibitor ([I]) and the combined cooperativity factor (αβ) by (λ):

$$CR - 1 = \frac{[I]\,(1 - \lambda)}{\lambda\,[I] + K_B} \qquad (4)$$

By multiplying with 1/λ we obtain the following equation:

$$CR - 1 = \left(\frac{1 - \lambda}{\lambda}\right)\left(\frac{[I]}{[I] + \frac{K_B}{\lambda}}\right) \qquad (5)$$

To better model the cooperative effects we further introduced the Hill-coefficient (n):

$$f([I]) = \left(\frac{1 - \lambda}{\lambda}\right)\left(\frac{[I]^n}{[I]^n + \left(\frac{K_B}{\lambda}\right)}\right) \qquad (7)$$

**Linear Schild Equation**

To fit the linear part of the Schild-plot, we adapted the Schild equation from literature describing agonist-antagonist affinity interactions at receptors.[2]

$$\log(CR - 1) = \log([B]) - \log(K_B) \qquad (8)$$

Due to the additional cooperative effect, besides the competitive the slope is deviating from 1, therefore we introduced the factor m, that describes the slope of the resulting curve. [B] is replaced by [I] for the inhibitor concentration.

$$\log(CR-1) = f([I]) = m \log([I]) - m \log(K_B) \qquad (9)$$
